# Activated SKN-1 alters the aging trajectories of long-lived *C. elegans* mutants

**DOI:** 10.1101/2024.07.09.602737

**Authors:** Chris D. Turner, Sean P. Curran

## Abstract

In the presence of stressful environments, the SKN-1 cytoprotective transcription factor is activated to induce the expression of gene targets that can restore homeostasis. However, chronic activation of SKN-1 results in diminished health and a reduction of lifespan. Here we demonstrate the necessity of modulating SKN-1 activity to maintain the longevity-promoting effects associated with genetic mutations that impair *daf-2*/insulin receptor signaling, the *eat-2* model of caloric restriction, and *glp-1*-dependent loss of germ cell proliferation. A hallmark of animals with constitutive SKN-1 activation is the age-dependent loss of somatic lipids and this phenotype is linked to a general reduction in survival in animals harboring the *skn-1gf* allele, but surprisingly, *daf-2lf; skn-1gf* double mutant animals do not redistribute somatic lipids which suggests the insulin signaling pathway functions downstream of SKN-1 in the maintenance of lipid distribution. As expected, the *eat-2lf* allele, which independently activates SKN-1, continues to display somatic lipid depletion in older ages with and without the *skn-1gf* activating mutation. In contrast, the presence of the skn-1gf allele does not lead to somatic lipid redistribution in *glp-1lf* animals that lack a proliferating germline. Taken together, these studies support a genetic model where SKN-1 activity is an important regulator of lipid mobilization in response to nutrient availability that fuels the developing germline by engaging the *daf-2/*insulin receptor pathway.

## INTRODUCTION

Since the discovery of insulin signaling mutants that nearly double the normal lifespan of worms, *C. elegans* has become an attractive model to understand longevity and aging^1-5^. The longevity of *C. elegans* can be manipulated via several pathways, including germline ablation, caloric restriction and deficiencies in insulin growth factor signaling^6,7^. Laser ablation of germline progenitor cells disrupts germline to soma signaling and promotes an increase in lifespan^8^. Null mutants for the *C. elegans* NOTCH orthologue glp-1, when raised at a restrictive temperature, will increase longevity via germ to soma signaling through pathways that modulate DAF-16 activity^9^. Broadly, dietary restriction is a driver of increased longevity^7,10,11^. In *C. elegans*, the nicotinic acetyl-choline receptor EAT-2 regulates pharyngeal muscle contractions regulating feeding rate^7^. Reduced feeding rate via *eat-2lf* mutation results in dietary restriction, several factors such as *pha-4* and *hlh-30* feed into *eat-2lf* mediated dietary restriction generally resulting in increased autophagy and proteasomal turnover, which promotes longevity^12,13^. Lastly, under nominal conditions, insulin-IGF1-like signaling opposes SKN-1 activation by signaling through to kinases such as SGK-1 which in turn phosphorylate SKN-1 and inhibit its nuclear accumulation ^14^. A reduction in insulin-IGF1-like signaling (IIS) causes an increase in SKN-1 activity that promotes stress resistance and longevity^15^.

SKN-1 is a multifaceted transcription factor that regulates the response to xenotoxins, unfolded proteins, and metabolic stress by upregulating appropriate stress response pathways to mitigate the stressful condition^14,16^. Loss of SKN-1 function results in a diminished stress response and a reduction in lifespan^14,16^. Although, mild overexpression of SKN-1, albeit without additional genetic or environmental activation was shown to increase lifespan^16^, investigations utilizing a genetically encoded constitutively activated *skn-1* mutation determined that longevity is severely diminished SKN-1 activity cannot be turned off^14,17^. Constitutively active *skn-1* mutants display a remarkable resistance to acute exposures to stress^17^, but at the cost of dysregulated lipid metabolism and mobilization^18^ that ultimately reduces overall lifespan^14,17,18^. This phenomenon is also observed in wild type animals exposed to pathogens^19^, but importantly represents a tradeoff between organismal stress responses and cytoprotection that influences survival that is mediated through lipid storage and distribution. Importantly, SKN-1 regulates cellular detoxification^14^, proteostasis^14^, and metabolism^14^ responses and receives input from multiple regulatory pathways; including nutrient sensation^14^, germline integrity^20^, and insulin signaling^15^, which when impaired can result in an extension of lifespan^14,16^.

Given the opposing longevity outcomes of SKN-1 activation in response to loss of *daf-2* signaling, germline proliferation, and pharyngeal pumping versus constitutive activation of SKN-1 from gain-of-function alleles, we sought to assess how constitutive SKN-1 activity would impact well-established longevity promoting mutations.

## RESULTS

### Constitutive activation of SKN-1 reduces the lifespan of canonically long-lived mutants

Animals harboring the gain-of-function (gf) *skn-1* allele, *skn-1(lax188)*^17^, display constitutive activation of the SKN-1 transcription factor that results in accelerated aging that manifests in a significant reduction of mean, median, and maximal lifespan^18,19,14^. To assess the impact that constitutive SKN-1 activation has on the longevity of canonically long-lived *C. elegans* mutants we generated double mutants of the *skn-1gf* allele and three long-lived mutants, *daf-2lf* ^2,21^, *eat-2lf* ^7,11^, or *glp-1lf* ^8,22^. We found that constitutive activation of SKN-1 shortened the lifespan of each of the long-lived mutants, but to varying magnitude (**Figure 1A-C**, Table S1). Specifically, the *eat-2lf;skn-1gf* mutant displays a median lifespan of 9 days most similar to the *skn-1gf* mutant alone and down from an extended 25-day median lifespan in the *eat-2lf* single mutants. Similarly, *glp-1lf;skn-1gf* double mutant animals display a median lifespan of 12 days that is down from 27 days for the *glp-1lf* single mutant. Finally, although the *daf-2lf;skn-1gf* double mutant animals display a significant reduction in median lifespan as compared to the *daf-2lf* mutants alone, the presence of the *daf-2lf* mutation did significantly increase lifespan in the context of the *skn-1gf* mutation; *daf-2lf;skn-1gf* double mutant animals are longer lived than WT animals (Table S1).

**FIGURE 1.**
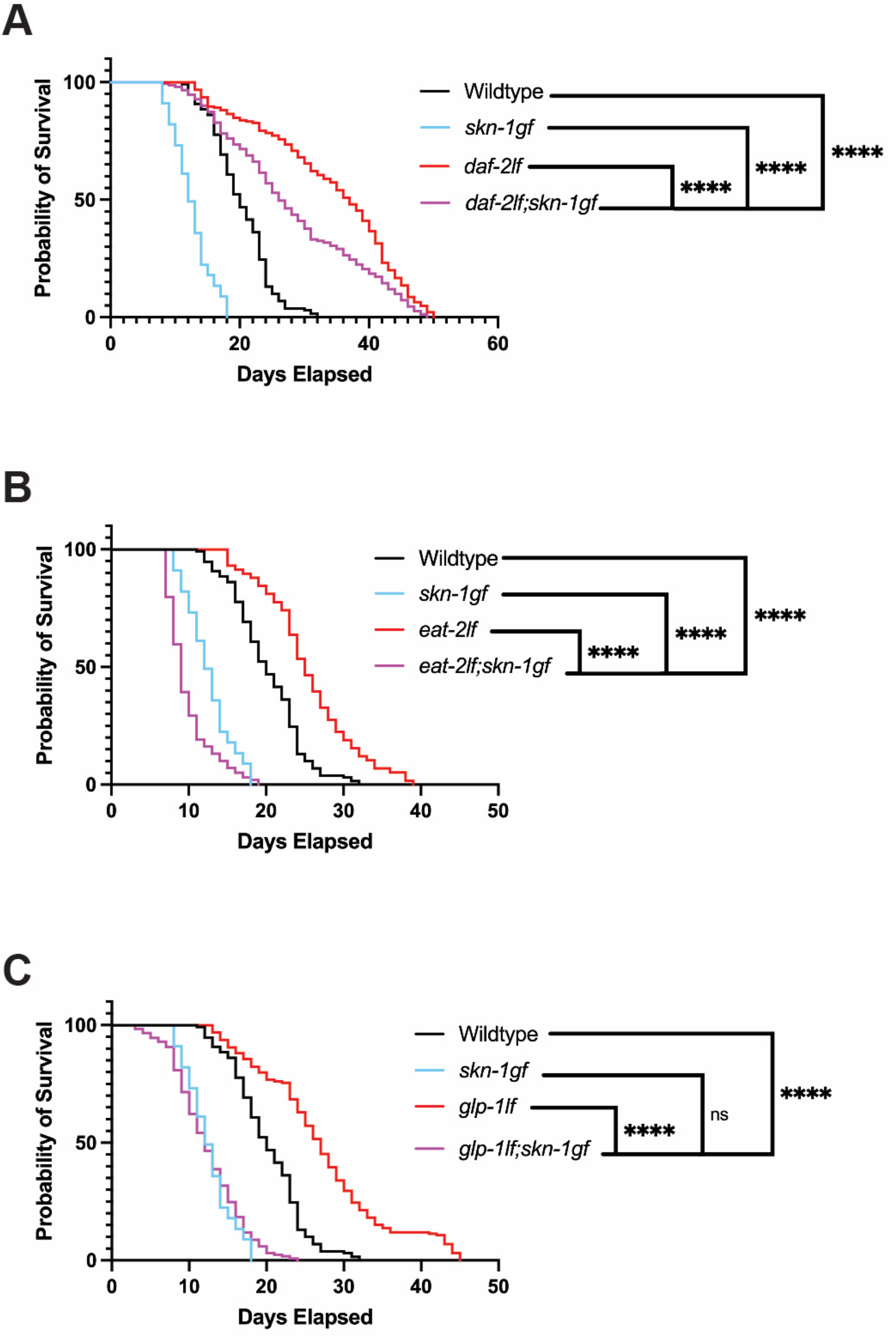
Constitutive SKN-1 activity alters lifespan trajectories. **(A)** Lifespan of *daf-2lf;skn-1gf* double mutant compared to wild type (WT), *daf-2lf*, and *skn-1gf*. **(B)** Lifespan of *eat-2lf;skn-1* double mutant compared to wild type (WT), *eat-2lf*, and *skn-1gf*. **(C)** Lifespan of *glp-1lf;skn-1* double mutant compared to wild type (WT), *glp-1lf*, and *skn-1gf* controls. Each condition (n=50; N=3) and analysis performed via long-rank Mantel-Cox test; ****, p<0.0001

Impaired IIS in *daf-2lf* mutants drives the alternative developmental program that leads to the stress resistant and long-lived dauer diapause state^23^. As such, we examined the influence of constitutive SKN-1 activation on dauer development by assessing the ability of *daf-2lf;skn-1gf* double mutant animals to enter the dauer diapause state at 25C and found no remarkable differences, as compared to the *daf-2lf* single mutant (Figure S1A). Similarly, the *daf-2lf;skn-1gf* mutants exited dauer diapause and continued development to adulthood when returned to the permissive temperature (Figure S1B). Taken together, these data suggest that the impact of constitutive SKN-1 transcriptional activity is specific to post-dauer decision activities mediated by the insulin signaling pathway.

### Impaired insulin signaling results in reduced lipid metabolism transcripts

Similar to the canonical negative regulation of the FoxO transcription factor DAF-16^24^, an established role of the IIS pathway is to restrict the activity of SKN-1^15^. However, in light of our finding that the *daf-2lf* mutation was capable of significantly increasing the lifespan of the constitutively activated *skn-1gf* mutant, we next examined the transcriptional changes that occur when animals harbor either *skn-1gf, daf-2lf*, or both alleles. Perhaps unsurprisingly, the transcriptomes to *daf-2lf and skn-1gf* single mutants were significantly different from each other and also WT animals (**Figure 2A**, Figure S2A-B, Table S2), but strikingly, the *daf-2lf; skn-1gf* was remarkably similar to the *daf-2lf* mutant alone (**Figure 2A**).

**FIGURE 2.**
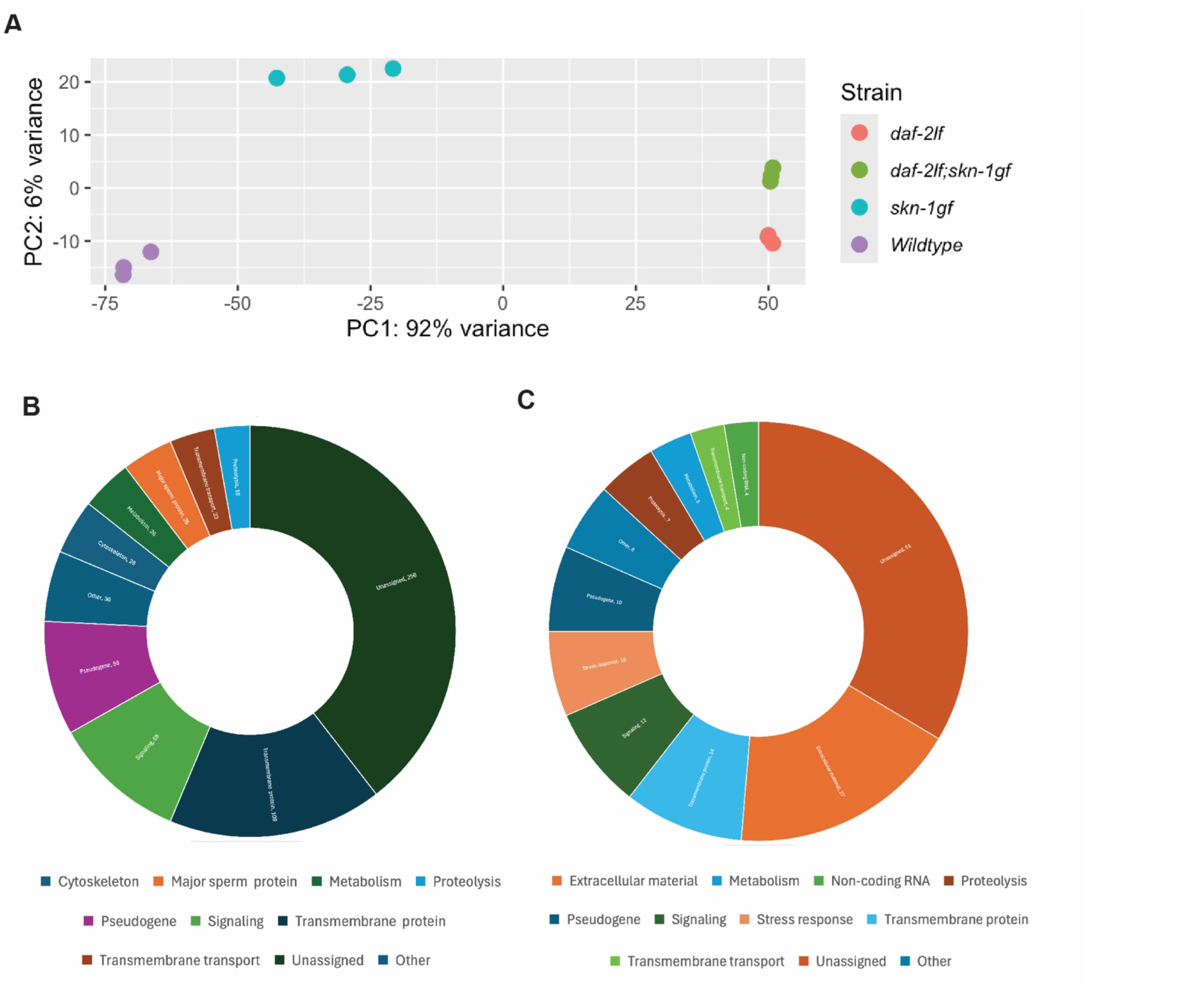
Loss of *daf-2* alter the *skn-1gf* transcriptional landscape. (**A**) PCA plot to determine group clustering, *daf-2lf* mutants cluster closely with *daf-2lf;skn-1gf* double mutants. (**B**) Sunburst chart of enriched GO terms found in down regulated genes from *daf-2lf;skn-1gf* vs *skn-1gf* comparison. (**C**) Sunburst chart of enriched GO terms found in the upregulated genes from *daf-2lf;skn-1gf* vs *skn-1gf* comparison. GO analysis performed using WormCat 2.0 using genes >4.5fold or <-7.5 fold to analyze the most differentially expressed genes, GO terms for all genes found in Table S2.

With the goal of trying to elucidate the molecular basis underlying the enhanced longevity effect of the *daf-2lf* allele on animals with constitutive SKN-1 activation, we focused our examination on the transcriptional differences between *skn-1gf* and *daf-2lf;skn-1gf* mutant animals that display a clear separation within the principal component space since the transcriptional signatures of the *daf-2lf* and *daf-2lf;skn-1gf* mutants animals were similar. We found 3487 genes were differentially regulated by at least 1.5-fold and adjusted p-value less than 0.05 (Figure S2B, Table S2). An analysis of GO-terms associated with the differentially effected genes revealed enrichment for signaling, specifically lipid signaling, and metabolism classes of genes (**Figure 2B-C**, Table S2). In addition, the majority (83%) of the top 100 differentially regulated genes in the *daf-2lf;skn-1gf* mutants animals, are previously confirmed targets of DAF-16^25^ (Table S3). Strikingly, the *daf-2lf;skn-1gf* double mutant animals display reduced expression of lipid metabolism genes in classes such as fatty acid chain elongation, beta oxidation, and acetyl transferases(Figure S2C, TableS2), which given the well-established connection between lipid homeostasis, SKN-1 activity, and lifespan^17,18,26,27^ could explain the increased longevity of *skn-1gf* mutants when IIS is impaired.

### Constitutive activation of SKN-1 alters age dependent somatic lipid redistribution in long-lived mutants

As previously reported, *skn-1gf* mutant animals display an age-dependent somatic depletion of fat (Asdf) phenotype where somatic lipid stores are depleted by mobilizing these lipids to the germline by vitellogenins^18,28^. As such, we used Oil-Red-O (ORO) staining to assess the distribution of intracellular lipid stores in the somatic and germline tissues of animals harboring the *daf-2lf* allele, *skn-1gf* allele, or both. We found that the loss of insulin signaling in *daf-2lf* mutant animals was capable of fully suppressing the Asdf phenotype observed in *skn-1gf* mutants (**Figure 3A**, Figure S3). We next examined the impact of the *eat-2lf* and separately the *glp-1lf* mutations on the *skn-1*-dependent somatic lipid depletion phenotype. In support of our previous findings^18^, loss of *eat-2*, which results in reduced food consumption^7^ that can independently activate SKN-1^29^, resulted in reduced somatic lipid stores (**Figure 3B**, Figure S3), but surprisingly, the *glp-1lf* mutation suppressed Asdf (**Figure 3C**, Figure S3), which suggests that somatic lipid depletion requires active proliferation of germ cells in the gonad.

Taken together, our genetic analysis of the impact that constitutive SKN-1 activity exerts on established genetic perturbations that drive longevity suggests an interaction model where *skn-1* pathways monitor nutrient uptake and integrate with the insulin signaling pathway in order to induce somatic lipid redistribution, which cannot occur without signals from the proliferative germline (**Figure 3D**).

**FIGURE 3.**
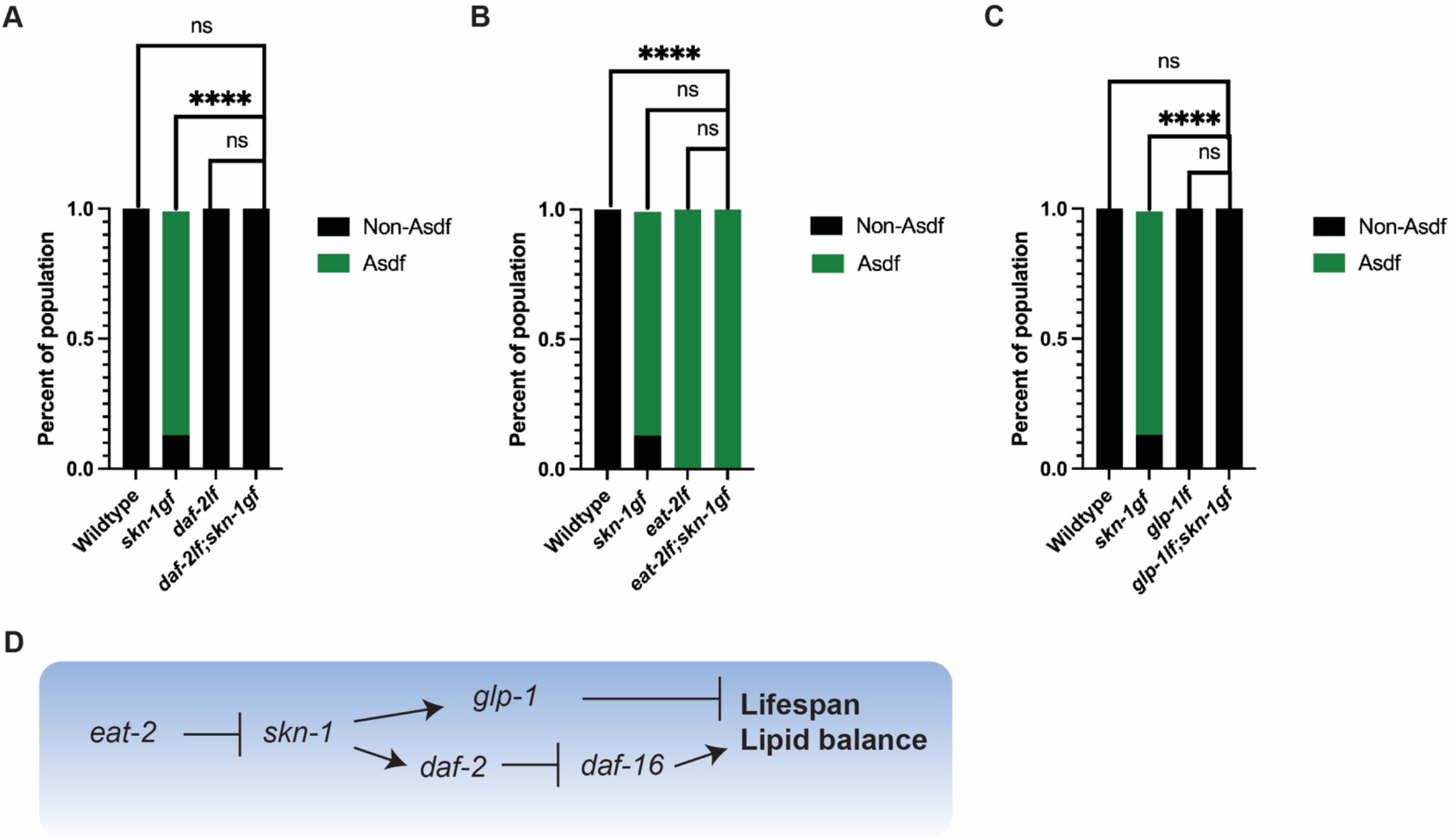
Age-dependent somatic depletion of fat is altered in long-lived mutants. **(A)** The *daf-2lf* allele suppresses Asdf in *skn-1gf* mutants. **(B)** The *eat-2lf* allele induces Asdf similar to *skn-1gf* mutants. **(C)** The glp-1lf allele suppresses Asdf in *skn-1gf* mutants. (**D**) Model of genetic relationship of longevity mutants in the context of constitutive SKN-1 activation. Each condition (n=100 N=3). Statistical comparisons made using a two-way ANOVA ****, p<0.0001

## DISCUSSION

The biology of aging research field has focused on single gene mutations that promote longevity and separately mutations that can accelerate aging, but interactions between the two remain understudied. The complexity of the aging process and the genetic variation that contributes to health outcomes requires an examination of effects of genetic variation in polygenetic models. The constitutively active SKN-1 mutants display accelerating aging phenotypes yet present an interesting paradox, where moderate activation ^16^ and even moderate overexpression^16^ of SKN-1 can be beneficial for the worm and even required in some circumstances to positively modulate lifespan however having SKN-1 always active results in a severe reduction in lifespan^14-17,29,30^. Our combination of constitutively active *skn-1* mutations with established longevity-promoting mutations has allowed us to uncover new ways that SKN-1 cytoprotection can interact with longevity paradigms.

While *eat-2lf* mutants are calorically restricted due to the reduction in feeding rate^7,31^, *skn-1gf* mutants replicate a starvation response by the induction of lipid utilization genes^17^. Our findings confirm that the *eat-2lf* mutants undergo somatic redistribution of lipids to the germline which is in line with the activation of SKN-1 that occurs in response to the actual starvation state of these mutants^18^. Furthermore, our observation that the *eat-2lf;skn-1gf* are exceptionally short-lived confirms previous observations that the degree of SKN-1 constitutive activation can correlate with the measured reduction in lifespan^17,32^. The previous observation that SKN-1 activity in the pair of ASI sensory neurons is required for longevity of dietarily restricted animals^29^ and for Asdf^32^ lends further evidence to the role(s) SKN-1 can play in monitoring nutrient status^29^.

The insulin/IGF-1 signaling pathway is an evolutionarily conserved modulator of healthspan and lifespan^21^. Although DAF-16/FoxO mediates the longevity responses through insulin signaling^33^, SKN-1 has also been demonstrated to potentiate the physiological outcomes from reduced insulin signaling^15^, but intriguingly our finding that impaired insulin signaling can increase the lifespan of animals with constitutively activated SKN-1 suggests that insulin-like signaling has roles downstream of constitutive SKN-1 activity. Alternatively, previous work has demonstrated that redirection of SKN-1 activity away from specific promoters can mitigate the negative health outcomes of constitutive SKN-1 activity^19^. It is possible that the activation of DAF-16 in the *daf-2lf* mutants could result in a competition model that redirects the transcriptional focus of SKN-1gf targets (**Figure 3D**) that results in the maintenance of somatic lipids and improves lifespan. Another way of analyzing the genetic relationship between *skn-1gf and daf-2lf*, is that rather than impaired insulin signaling being beneficial to *skn-1gf*, constitutively activated SKN-1 is detrimental to animals with impaired insulin signaling. Given that SKN-1 activation can suppress DAF-16 mediated stress tolerance^34^ it is possible that constitutive SKN-1 activation abrogates the longevity promoting effects of DAF-16 in insulin signaling mutants. This idea is supported by our transcriptomic analysis, where we find that the expression of several DAF-16 target genes are altered in the *daf-2lf; skn-1gf* animals as compared to *daf-2lf* mutants alone. Furthermore, our examination of lipid homeostasis has provided insight into the role of insulin signaling plays in the distribution of stored intracellular lipids. *daf-2lf* mutants have been demonstrated to store increased levels of lipids in their somatic tissues ^35^ and our findings reveal that even with constitutive activation of SKN-1, *daf-2lf* mutant animals do not deplete somatic lipids as *skn-1gf* mutants. This suggests that SKN-1, when activated, requires the insulin signaling pathway to modulate lipid redistribution to the germline.

The relationship between reproduction and organismal longevity across organisms is well-established^36-38^. In *C. elegans*, the specific loss of germline progenitor cells leads to a significant increase in lifespan^19^ and similarly, the loss of the germ cell proliferation mediated by GLP-1/NOTCH signaling can extend lifespan^5^. The finding that *glp-1lf;skn-1gf* double mutant lifespan was unremarkable as compared to *skn-1gf* single mutants suggests that the GLP-1/NOTCH signaling effect on longevity requires the ability to turn off SKN-1 activation. Interestingly, *glp-1lf;skn-1gf* double mutants do not undergo redistribution of somatic lipids which uncouples reduced lifespan and Asdf phenotypes associated with constitutive SKN-1 activity. This finding also suggests that a germ cell proliferation is required for lipid redistribution and not simply the somatic gonad, which *glp-1lf* mutants still possess. This could indicate that during stressful events that activate SKN-1, the proliferative germline signals to somatic tissues like the intestine where lipids are stored, to mobilize these resources to promote reproduction and ensure fitness^18^.

Collectively, our study demonstrates the need to tight regulate SKN-1 activity in the context of long-lived genetic background. Although other genetic pathways are likely influenced by constitutive SKN-1 activity our work provides a model to approach the examination of the polygenic nature of organismal longevity and health across the lifespan.

## SUPPLEMENTARY FIGURES

**Figure S1.**
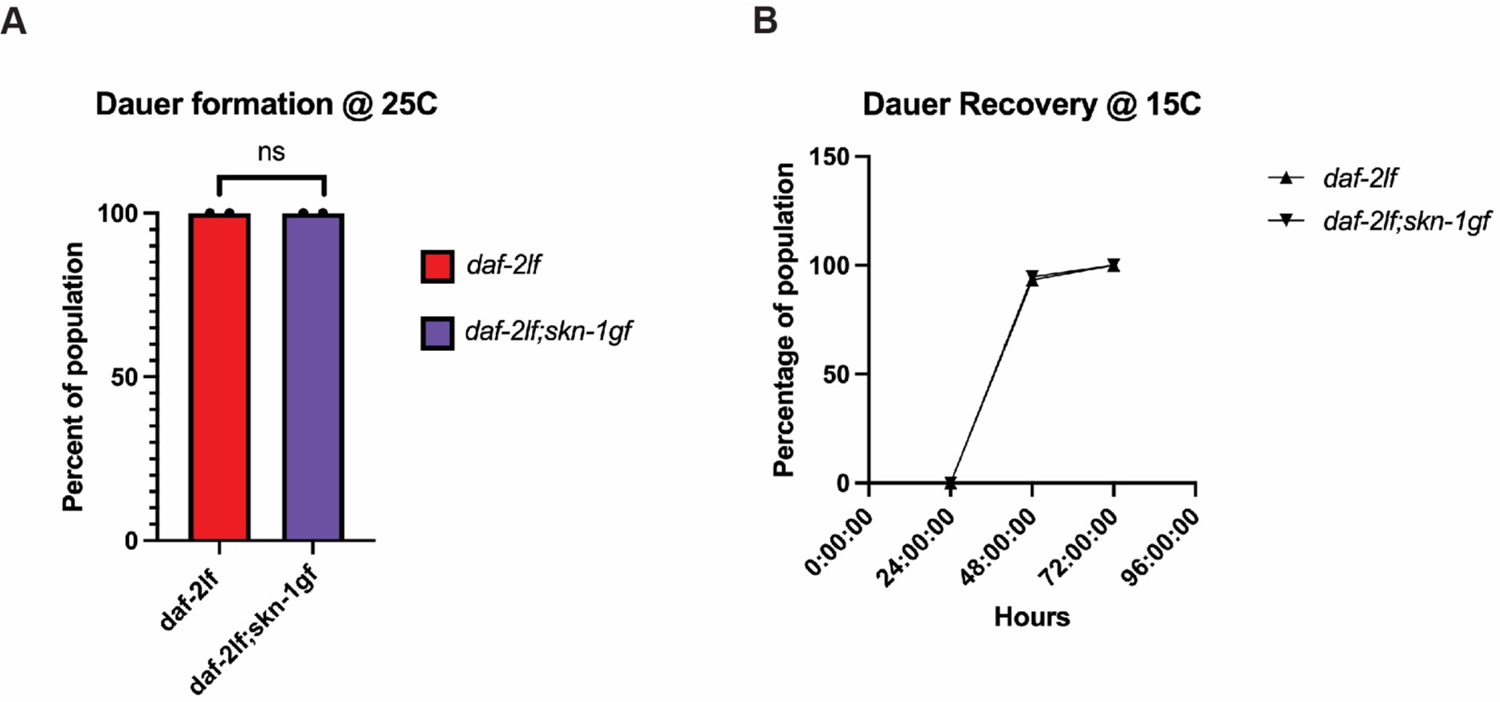
Constitutively activated SKN-1 does not impact daf-2lf dauer entry and exit. Dauer formation (**A**) and recovery (**B**) of *daf-2lf* mutants at 25C is unaffected by the *skn-1gf* allele.

**Figure S2.**
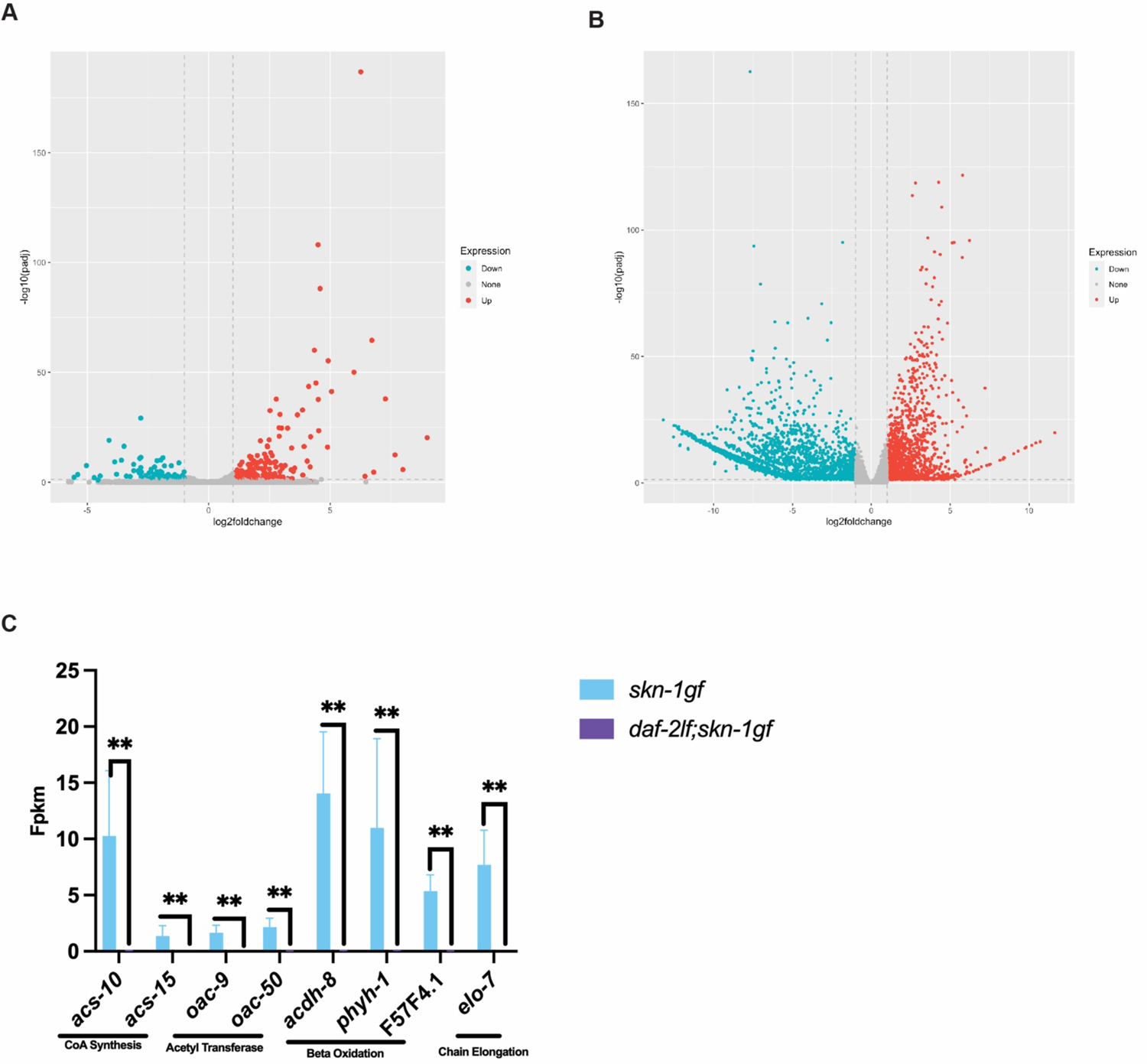
Transcriptional analysis of daf-2lf;skn-1gf double mutant. (**A**) Volcano plot comparing *daf-2lf* and *daf-2lf;skn-1gf* differentially expressed genes. **(B)** Volcano plot comparing *daf-2lf;skn-1gf* double mutants to *skn-1gf* mutants. **(C)** Decreased expression of lipid metabolism genes in the *daf-2lf;skn-1gf* double mutant.

**Figure S3.**
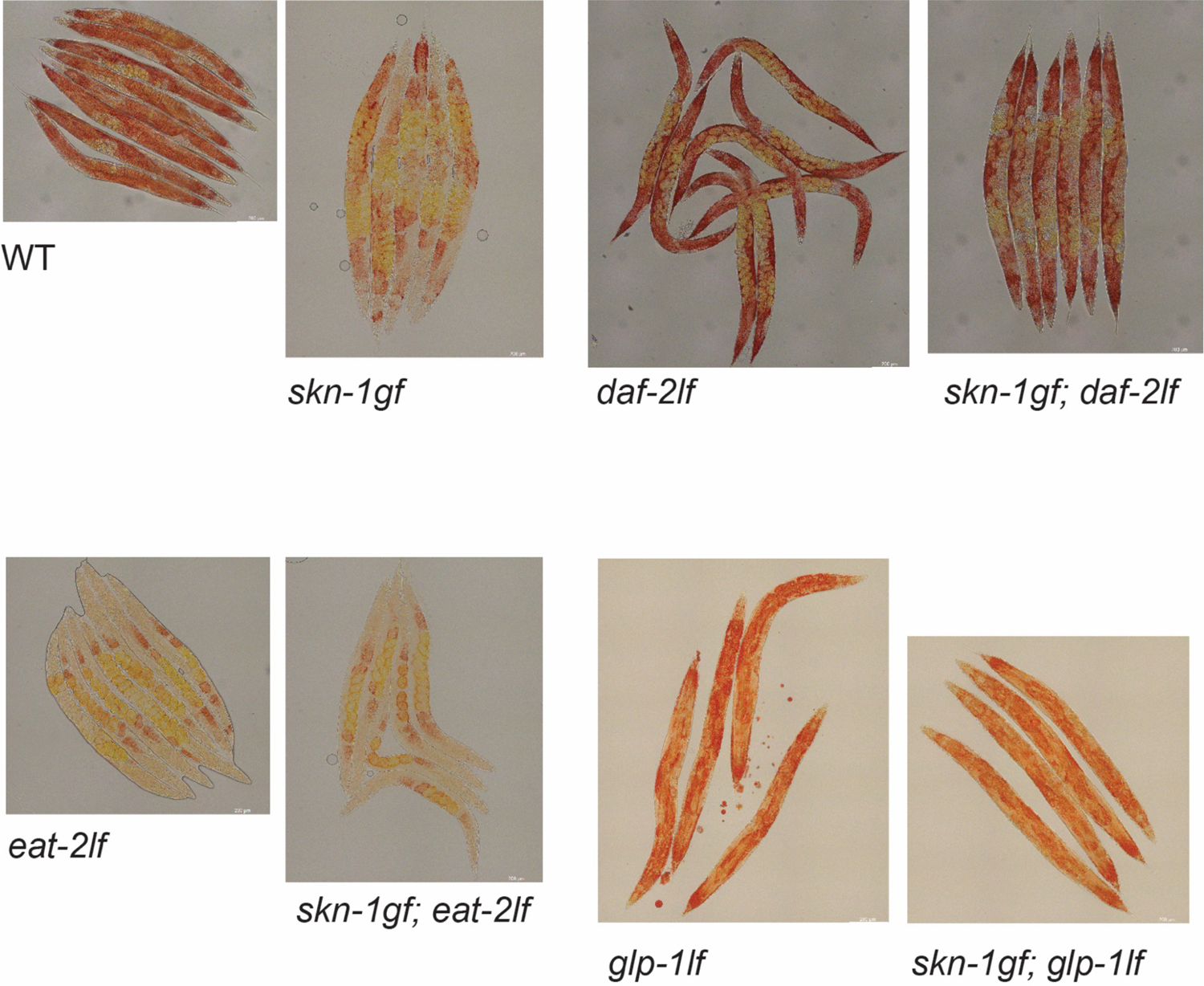
ORO staining of longevity mutants with constitutive SKN-1 activation. Representative images of Oil-red-O (ORO) staining of lipid droplets in animals of the indicated genotypes.

## SUPPLEMENTARY TABLES

Table S1. Lifespan data

Table S2. RNAseq

Table S3. Overlap of DAF-16 regulated genes with top 100 differentially regulated genes in the *daf-2lf;skn-1gf* mutants animals

## ACKNOWLEDGMENTS

We thank S. Ledgerwood for technical assistance. This work was funded by the NIH R01AG058610 and Hevolution Foundation award HF AGE-004 to SPC, and T32AG052374 to CDT. We also thank the USC School of Gerontology Imaging Core that is funded in part by the Nathan Shock Center of Excellence P30AG068345. Some strains were provided by the CGC, which is funded by the NIH Office of Research Infrastructure Programs (P40 OD010440). We thank WormBase for database curation and data access.

## Author contributions

Conceptualization: SPC; Methodology: SPC; Investigation: CDT and SPC; Visualization: CDT and SPC; Supervision: SPC; Writing: CDT and SPC.

## Competing interests

All authors declare that they have no competing interests.

## Data availability

Plasmids and strains available upon request. All data are available in the main text or the supplementary materials. RNAseq data is available at the NIH (GEO) Gene Expression Omnibus GSE270810

## MATERIALS AND METHODS

### *C.elegans* strains and maintenance

*C.elegans* were raised on 6cm nematode growth media(NGM) plates supplemented with streptomycin and seeded with OP50. All worm strains were grown at 20°C except for temperature sensitive strains which were grown at 15°C, worm strains were unstarved for at least three generations before being used^39^.

Strains used in this study: WT, N2 Bristol; SPC227, *skn-1(lax188)*; DA456, *eat-2(ad456)*; CB1370, *daf-2(e1370)*; CB4037, *glp-1(e2141)*.

Double mutants used in this study were obtained by standard genetic techniques. Some strains used in this study were provided by the Caenorhabditis Genetics Center, which is funded by the NIH Office of Research Infrastructure Programs (P40 OD0104400).

### Lifespan Analysis

Synchronized L4 animals were moved onto NGM plates seeded with OP50 without the supplementation of FuDR. All worms were kept at 20°C and were transferred each day during the reproductive period. Worms that died of vulval bursting, bagging, or crawling off the plate were censored. Lifespan analysis and graphing was performed using GraphPad Prism 10.

### Dauer Entry and Exit Assay

Gravid adult worms were bleach spotted and progeny were raised at 25°C for 30 hours (hrs) to allow for the development of dauer larvae. Dauer were counted and proportions calculated as percent of total population. For the exit assay, dauers were shifted to 15°C and plates were scored for the presence of non dauers at 24 hrs, 48 hrs, and 72 hrs.

### RNA-seq Analysis

Analysis performed as in Turner et al. 2023^32^, in brief gravid adult worms were egg prepped and eggs were allowed to hatch overnight for a synchronous L1 population. The next day, L1s were dropped on to seeded NGM plates and allowed to grow 48 hrs, 72 hrs, 120 hrs or 168 hrs (L4, Day 1 Adult, Day 3 Adult , Day 5 Adult) before collection. Animals were washed three times with M9 buffer and frozen in TRI reagent at -80°C until use. Animals were homogenized and RNA extraction was performed via the Zymo Direct-zol RNA Miniprep kit (Cat. #R2052). Qubit™ RNA BR Assay Kit was used to determine RNA concentration. The RNA samples were sequenced and read counts were reported by Novogene. Read counts were then used for differential expression (DE) analysis using the R package DESeq2 created using R version 3.5.2. Statistically significant genes were chosen based on the adjust p-values that were calculated with the DESeq2 package. Gene ontology analysis was performed using WormCat 2.0^40^.

### Oil Red O (ORO) Staining

Performed as in Stuhr et al. 2022^41^, in brief, gravid adult worms were egg prepped and allowed to hatch overnight for a synchronous L1 population. The next day, worms were dropped onto plates seeded with bacteria and raised to 120 hrs (Day 3 Adult stage). Worms were washed off plates with PBS+triton, then rocked for 3 min in 40% isopropyl alcohol before being pelleted and treated with ORO in diH2O for 2 hrs. Worms were pelleted after 2 hrs and washed in PBS+triton for 30 min before being imaged at 20x magnification with LAS X software and Leica Thunder Imager flexacam C3 color camera.

### Age-dependent Somatic Depletion of Fat (Asdf) Quantification

Performed as in Turner et al. 2023^32^, in brief, ORO-stained worms were placed on glass slides and a coverslip was placed over the sample. Worms were scored and images were taken with LAS X software and Leica Thunder Imager flexacam C3 color camera. Fat levels of worms were placed into two categories: non-Asdf and Asdf. Non-Asdf worms display no loss of fat and are stained dark red throughout most of the body (somatic and germ cells). Asdf worms had most, if not all, observable somatic fat deposits depleted (germ cells only) or significant fat loss from the somatic tissues with portions of the intestine being clear (somatic < germ).

